# Therapeutic acetaminophen exposure does not measurably alter human cortical organoid development

**DOI:** 10.1101/2025.10.07.681041

**Authors:** Luca Trovò, George Kucera, Gimena Gomez, Janaina Sena de Souza, Alysson R. Muotri, Lilia M. Iakoucheva

**Author notes:** Equal contribution (George Kucera, Gimena Gomez). Corresponding Authors (L.M.I) and (A.R.M).

## Abstract

Acetaminophen (APAP) is the most widely used analgesic during pregnancy, yet its effects on prenatal human brain development remain incompletely understood. Epidemiological studies have reported inconsistent associations between prenatal APAP exposure and later neurodevelopmental outcomes, underscoring the need for mechanistic evaluation in human-relevant developmental models. Here, we examined how APAP influences early cortical development using induced pluripotent stem cell-derived cortical organoids (COs) generated from six independent lines. Organoids were exposed to physiologically relevant APAP concentrations (25, 50, and 100 μM) for 5 days beginning at day 21 of differentiation, corresponding to late first-trimester cortical development. We assessed organoid growth, apoptosis, differentiation, synaptic maturation, transcriptomic profiles using bulk and single-nucleus RNA sequencing (snRNA-seq), and functional network activity using multielectrode array recordings up to 4 months. APAP exposure did not affect organoid size, cytoarchitecture, or viability. Neuronal and progenitor cell proportions, as well as synaptic puncta density were unchanged. Bulk RNA-seq revealed subtle transcriptional changes only at the highest dose (16 differentially expressed genes at 100 μM), enriched for neurodevelopmental pathways. In contrast, snRNA-seq at 3 months revealed no changes in cell type composition or gene expression. Consistent with these findings, electrophysiological measures including firing rate, burst frequency, and network synchrony, were indistinguishable from controls. Together, these results indicate that exposure to therapeutic APAP concentrations during a critical window of early cortical development produces minimal molecular perturbations without detectable consequences. This study provides mechanistic evidence indicating that, in this human organoid model, recommended APAP exposure is not associated with detectable disruptions in pathways implicated in brain development.

## INTRODUCTION

Acetaminophen (APAP) is the most commonly used analgesic and antipyretic during pregnancy, reported in over half of women worldwide^1,2^. Its popularity reflects a favorable safety profile: unlike Non-Steroidal Anti-Inflammatory Drugs (NSAIDs), it does not increase gastrointestinal or cardiovascular risks, and unlike opioids, it carries no risk of dependence or respiratory depression^3,4^. For decades, APAP has therefore been considered the drug of choice for managing pain and fever in pregnancy.

Concerns have increased, however, because APAP easily crosses the placenta and enters the fetal brain^5,6^. Over the past decade, epidemiologic studies have reported associations between prenatal APAP exposure and neurodevelopmental disorders (NDDs) including Autism Spectrum Disorder (ASD) and Attention-Deficit Hyperactivity Disorder (ADHD)^7–11^. Several studies suggest exposure-response relationships, especially with long-term use or exposures during the late first and second trimesters^9,12^. However, these findings remain controversial. Observational studies are susceptible to indication bias, because maternal conditions that prompt APAP use, such as fever and infection, may themselves increase ASD risk^13,14^. Additional concerns include residual confounding^15^ and exposure misclassification for over-the-counter medication use, particularly when ascertainment relies on maternal recall^16^.

Sibling-controlled analyses have helped address these limitations. In the Norwegian Mother and Child Cohort, associations observed in conventional models were no longer evident in within-family analyses^17^. Similarly, a Swedish population-based study of 2.5 million children and a recent Taiwanese cohort study of more than 2 million births both reported full-cohort associations with NDDs that were substantially reduced and no longer statistically significant in sibling-matched analyses ^18,19^.

Animal studies have raised concerns about potential long-term behavioral and synaptic consequences of developmental APAP exposure, with proposed mechanisms including oxidative stress, apoptosis, and endocannabinoid signaling^20–22^. However, many of these studies used supra-therapeutic doses, and species-specific differences in brain development and drug metabolism limit translation to humans^23^. In contrast, multiple *in vitro* studies in rodent neurons^24,25^ and human iPSC-derived neuronal and organoid models^26,27^, where acetaminophen has been used as a negative control, report no detectable effects of therapeutic-range exposure on neuronal differentiation or network activity within these systems. Together, these observations underscore a key unresolved question: whether acetaminophen, at concentrations relevant to therapeutic maternal use, directly perturbs early human cortical development.

Human induced pluripotent stem cell (hiPSC)-derived cortical organoids (COs) provide an experimentally tractable system for investigating early human cortical development *in vitro*. Given convergent genetic, transcriptomic, and developmental evidence implicating the cerebral cortex as a primary region of vulnerability in neurodevelopmental disorders^28–35^, we focused specifically on cortical development rather than modeling whole-brain effects. COs recapitulate fundamental features of fetal cortical development, including progenitor proliferation, neuronal differentiation, and progressive network maturation, and their transcriptional trajectories can be aligned with *in vivo* developmental timelines^36–38^. Consistent with this relevance, COs have been widely used to model both genetic (e.g., CHD8, 16p11.2 copy number variation) and environmental (e.g., alcohol, pesticide exposure) risk factors associated with ASD^39–41^.

In this study, we used COs to examine the effects of APAP exposure at physiologically relevant concentrations during a late first-trimester-equivalent developmental window. We evaluated organoid growth, apoptosis, cellular composition, transcriptomic profiles, and emergent network activity. Across these complementary readouts, APAP exposure produced only modest transcriptional changes and no detectable structural or functional consequences within this experimental system. These findings provide experimental context for recent familial-control epidemiologic studies^18^ while remaining limited to the scope of the *in vitro* model.

## RESULTS

### Cortical organoids maintain cytoarchitecture following APAP exposure

To investigate the potential neurodevelopmental effects of APAP exposure, we used hiPSC-derived COs (**Fig. 1A**) which recapitulate key features of early human brain development^36–38^. We initiated APAP exposure on day 21, corresponding to the transition from late first-to early second-trimester development based on the developmental mapping of our cortical organoid protocol^36^. This interval overlaps with a critical window of vulnerability to NDDs^28–35,42^. APAP exposure was administered for 5 consecutive days, reflecting the typical maximum duration of short-term clinical use of Acetaminophen during pregnancy.

**Figure 1.**
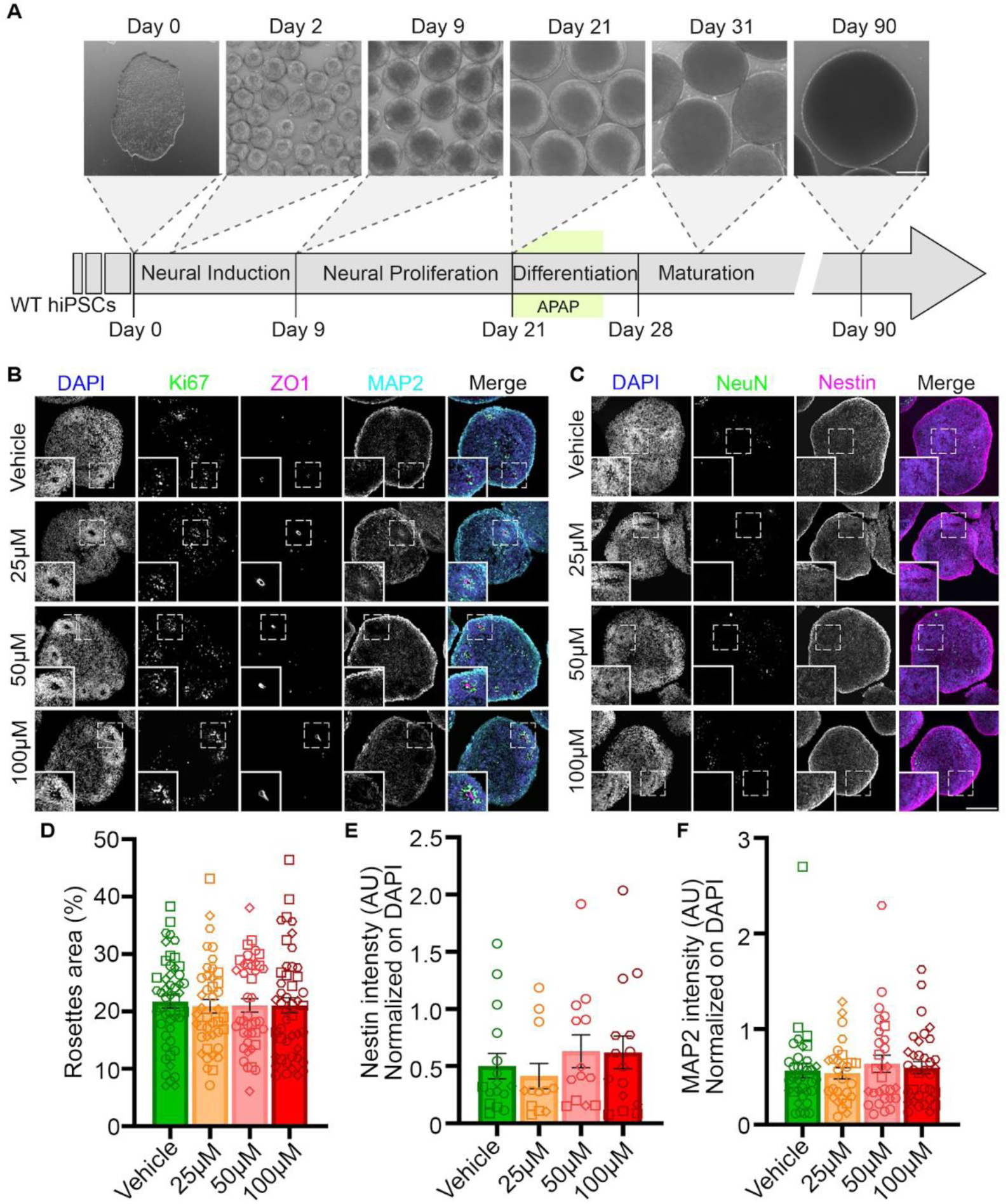
Neural rosette organization and cellular architecture remain intact across APAP exposure concentrations. **(A)** Schematic showing key developmental stages of COs: neural induction (days 0-9), neural proliferation (days 9-21), differentiation (days 21-28), and maturation (days 28+). Beginning on day 21, COs were exposed to the vehicle (PBS) or APAP (25 μM, 50 μM, or 100 μM) for 5 consecutive days. Scale bar: 200 μm. (**B-C).** Representative confocal images of 20 μm cryosections of one-month-old organoids immunostained for DAPI (blue), Ki67 (green), ZO1 (magenta), and MAP2 (cyan) **(B)** or DAPI (blue), NeuN (green), and Nestin (magenta) **(C).** Insets show higher magnification views of neural rosettes. Scale bars: 100μm (insets); 200μm (full images). (**D**) Quantification of neural rosette area expressed as percentage of total organoid area across vehicle, 25 µM, 50 µM, and 100 µM APAP conditions (n≥20 organoids, 4 iPSC lines, 4 independent batches). (**E-F**) Quantification of Nestin (E) and MAP2 (F) immunofluorescence intensity normalized to DAPI. Bars represent mean ± SEM. Each data point represents an individual organoid; different symbols indicate distinct iPSC lines (n≥10 organoids, 4 iPSC lines, 3 independent batches). Statistical significance was assessed using a mixed-effects model (REML) with Dunnett’s multiple comparisons test versus vehicle control. No significant differences were detected across exposure conditions.

Dosages (25 μM, 50 μM, and 100 μM) were selected based on physiologically relevant exposure levels. Prior *in vivo* studies in adults^43,44^ determined 100 μM as the peak maternal plasma concentration, while i*n silico* models of APAP transfer across the umbilical cord^45^ estimated 25 μM as the mean fetal arterial concentration, which we used as the lowest dose. This range enabled evaluation of dose-dependent effects spanning physiologically relevant fetal exposures (25 μM) to maternal therapeutic peak levels (100 μM).

To confirm that APAP exposure did not disrupt the cytoarchitecture of COs, we performed immunofluorescence (IF) quality control assessments focusing on neural rosette organization, a hallmark of CO development^37,38^. Across all conditions, COs developed well-organized neural rosette structures resembling the ventricular zone architecture of the developing human brain (**Fig. 1B**). These structures displayed characteristic features of proper neurodevelopment: ZO1-positive apical domains forming tight junction belts indicative of proper epithelial polarity; Ki67^+^ proliferative zones indicative of active neurogenesis; and MAP2^+^ neuronal processes surrounding the rosettes (**Fig. 1B**). NeuN^+^ post-mitotic neurons were appropriately distributed outside proliferative zones, while Nestin^+^ progenitor cells spread outward from apical domains, consistent with ongoing neurogenesis and preserved developmental progression (**Fig. 1C**). Quantification of neural rosette area (**Fig. 1D**), Nestin⁺ (**Fig. 1E**) and MAP2⁺ (**Fig. 1F**) revealed no significant differences across APAP exposure conditions. Together, these findings demonstrate that APAP exposure did not significantly disrupt organoid cytoarchitecture or neural organization.

### Acetaminophen exposure does not affect cortical organoid size, growth trajectories, or induce apoptosis

Neurodevelopmental disorders are often associated with atypical brain growth patterns, including micro-and macrocephaly^46,47^, and COs have been shown to recapitulate these phenotypes through measurable size differences^36^. To assess whether APAP exposure affects COs size and growth, we performed a longitudinal morphological analysis from the initiation of exposure (day 21) through day 31, and again at 3 months (**Fig. 2A**). COs were imaged every other day during the exposure period, and subsequently at 3-month time point.

**Figure 2.**
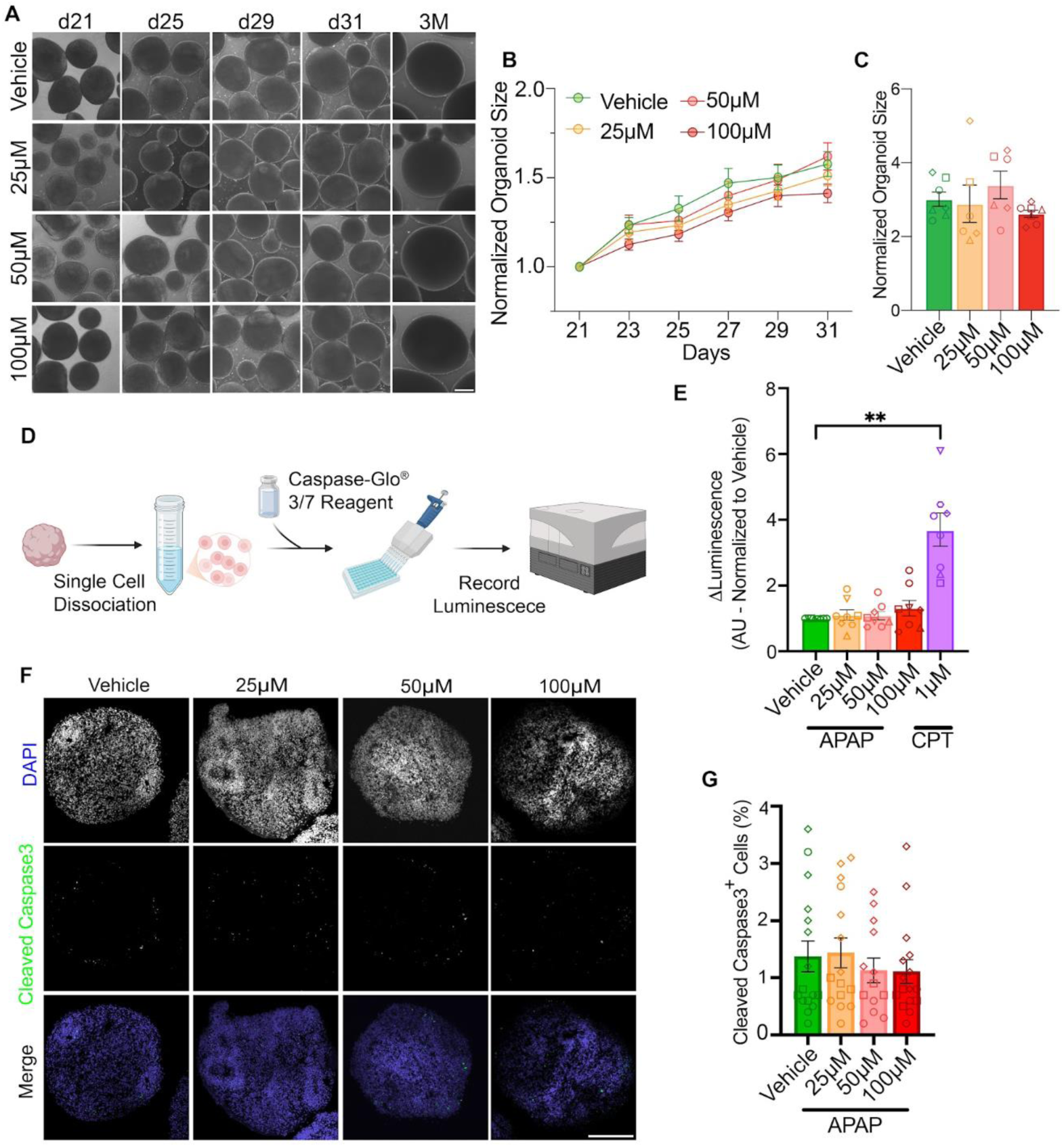
APAP exposure does not affect cortical organoid growth or induce apoptosis in cortical organoids. **(A)** Representative brightfield images of human iPSC-derived brain cortical organoids at sequential timepoints (days 21, 23, 25, 27, 29, 31, and 3 months) across APAP conditions (vehicle, 25 μM, 50 μM, 100 μM). **(B)** Longitudinal growth curves with areas normalized to day 21 (mean ± SEM) (n ≥1000 organoids, 6 iPSC lines, 4 independent batches). **(C)** Long-term organoid size at 3 months, normalized to day 21. Data represent mean ± SEM across batches, with different shapes indicating individual cell lines (n >100 organoids, 5 iPSC lines, 4 independent batches). Statistical analysis: mixed-effects model (REML) with Dunnett’s multiple comparisons test versus vehicle control. Scale bar: 200μm. **(D)** Schematic of the Caspase-Glo 3/7 luminescence assay workflow. Organoids were dissociated, mixed with Caspase-Glo® 3/7 Reagent, plated, and luminescence was measured by plate reader. **(E)** Quantification of caspase-3/7 activity (Δ-luminescence normalized to vehicle). CPT exposure served as a positive control. Data represent mean ± SEM (n=6 iPSC lines, 5 independent batches, 3 technical replicates per point). Statistical analysis: one-way ANOVA with Dunnett’s multiple comparisons (**p < 0.01; all APAP conditions p > 0.05 vs. vehicle). **(F)** Representative confocal images of 20μm cryosections from one-month-old organoids immunostained for DAPI (blue) and cleaved caspase-3 (green) across exposure conditions. Scale bar: 200 μm. **(G)** Quantification of cleaved caspase-3^+^ cells as a percentage of total cells. Data points represent individual organoids, while different shapes represent individual cell lines. Data presented as mean ± SEM (n≥10 organoids, 3 iPSC lines, 2 independent batches). Statistical analysis: mixed-effects model (REML) with Dunnett’s multiple comparisons test versus vehicle control. No significant differences were detected across exposure conditions.

Across all APAP concentrations, COs exhibited highly similar growth patterns (**Fig. 2B**). Normalized growth curves from day 21 to day 31 were nearly overlapping across exposure conditions, and long-term measurements revealed no significant differences in final organoid size at 3 months (**Fig. 2C**). Although a modest trend towards reduced size was observed at the highest dose (100 μM) during the exposure period, this effect did not reach statistical significance. Together, these results suggest that APAP exposure does not measurably impact organoid size or growth trajectories in this model.

To assess whether APAP exposure induces cytotoxicity, we performed two complementary assays. First, caspase-3/7 activity was measured in dissociated organoids at day 26, immediately following exposure, using the Caspase-Glo 3/7 luminescence assay (**Fig. 2D**), which quantifies apoptotic signaling. In parallel, organoids were treated with camptothecin (CPT; 1 μM for 18 h) as a positive control due to its well-established pro-apoptotic effects. APAP exposure did not increase caspase-3/7 activity relative to vehicle controls (**Fig. 2E**), whereas CPT treatment robustly activated caspase-3/7, confirming assay sensitivity. These results indicate that, under the conditions tested, APAP exposure did not significantly increase apoptosis in this cortical organoid model, enabling assessment of developmental effects in the absence of overt cytotoxicity.

Second, one-month-old organoid slices were stained for cleaved caspase-3, a marker of apoptosis (**Fig. 2F**). In all conditions, apoptotic cells remained below 1% of the total cell population, with no significant differences between APAP-treated samples and vehicle controls (**Fig. 2G**). These consistently low apoptosis levels are typical of healthy organoid cultures and suggest there is no increase in apoptotic activity. Altogether, these results indicate that APAP exposure at the tested doses did not significantly induce apoptosis or cytotoxicity in our CO model.

### Acetaminophen exposure causes subtle dose-dependent transcriptional changes that impact neurodevelopmental pathways

Having established that APAP exposure does not cause changes in cytoarchitecture and does not induce apoptosis, we next examined whether it produces more subtle molecular effects using bulk RNA sequencing of 48 samples (4 conditions: control, 25, 50, and 100 µM APAP × 6 lines × 2 technical replicates), with replicates generated from independent organoid batches grown on different days. After rigorous quality control and detection of sample outliers (**Fig. S1**), we performed differential gene expression analysis with a limma-voom model, incorporating a duplicate correlation to account for replicates. Despite detecting a relatively small number of differentially expressed genes (DEGs; 0-16), we observed a clear dose-dependent response with no significant DEGs at 25 μM, ten at 50 μM, and sixteen at 100 μM using thresholds of |log2 fold change| ≥ 0.2 and FDR-adjusted p < 0.05 (**Fig. 3A-C, Supplementary Table S1**). This trend indicates that APAP exposure causes subtle transcriptional changes in a concentration-dependent manner.

**Figure 3.**
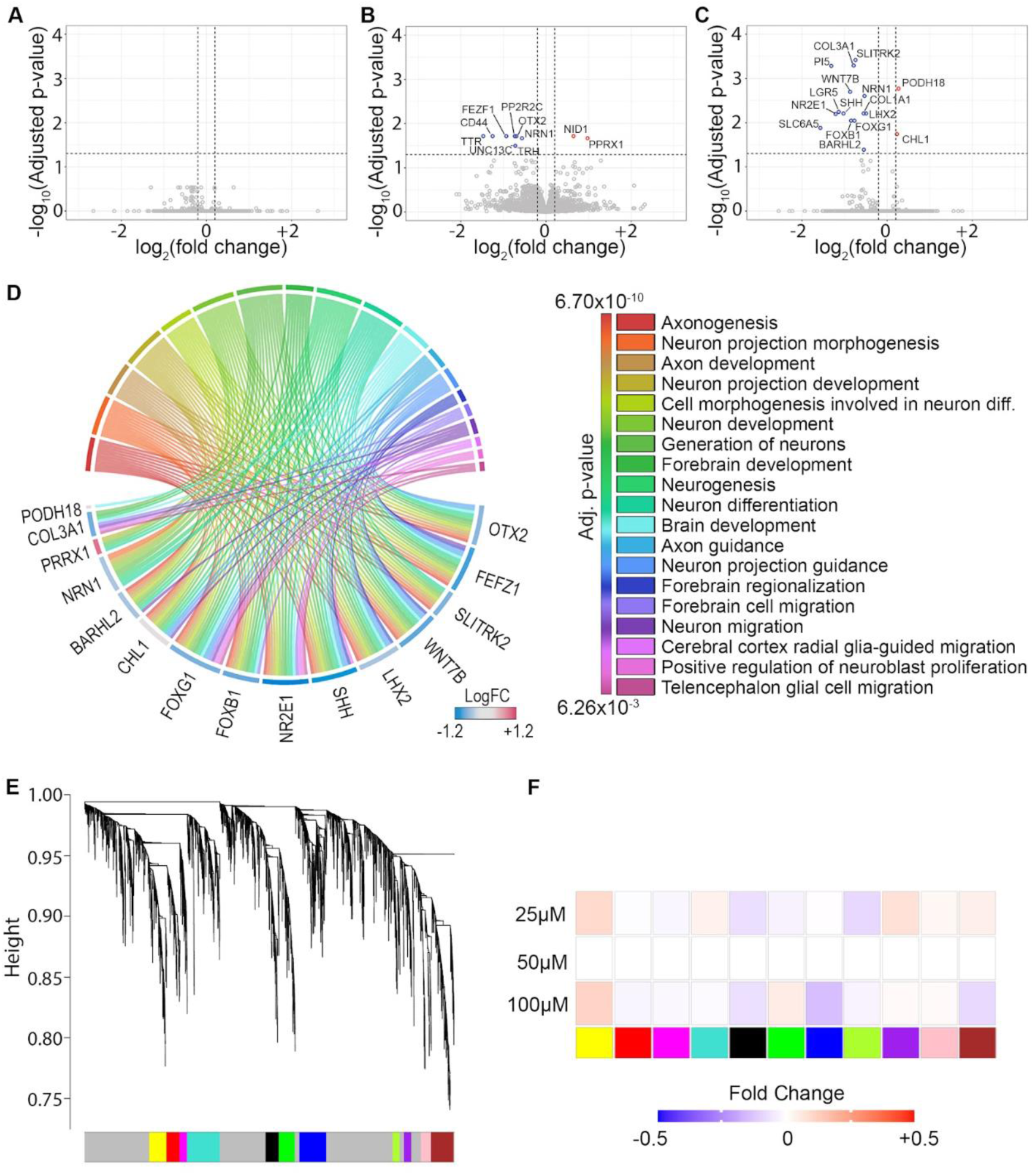
Dose-dependent transcriptional changes in cortical organoids after APAP exposure. **(A-C)** Volcano plots showing differentially expressed genes for 25 μM (**A**), 50 μM (**B**), and 100 μM (**C**) APAP *vs.* vehicle control. Genes meeting the differential expression criteria (|log2 fold change| ≥ 0.2 and FDR-adjusted p < 0.05) are shown in red (upregulated) or blue (downregulated), whereas non-significant genes are shown in gray (n=5 lines, 2 technical replicates per line, adjusted p<0.05). (**D)** Chord diagram illustrating associations between significantly dysregulated genes and enriched Gene Ontology (GO) biological processes related to neural development. **(E)** Dendrogram from WGCNA showing hierarchical clustering of co-expressed gene modules. **(F)** Module-exposure relationship heatmap showing the average fold change of module eigengenes in response to APAP exposure (25 µM, 50 µM, and 100 µM) relative to vehicle control. Warmer colors indicate positive correlations (upregulation), cooler colors indicate negative correlations (downregulation) of module eigengenes. No modules showed significant association with APAP dose using Benjamini–Hochberg FDR correction.

Given the inherent variability of organoid systems, we next assessed whether the direction of fold-change for each DEG was shared across hiPSC lines. Most DEGs showed the same direction of effect in at least 60% of lines (**Fig. S2**), indicating that the observed transcriptional changes were concordant across multiple lines, and no DEG was driven exclusively by a single hiPSC line. To assess the functional impact of these DEGs, we performed Gene Ontology (GO) enrichment analysis on genes identified at 50 μM and 100 μM conditions (**Supplementary Table S2**). The chord diagram (**Fig. 3D**) shows an overrepresentation of neurodevelopment-related GO terms, including early neurogenesis, brain patterning, and neuronal development.

To investigate whether APAP influences higher-order transcriptional changes, we carried out Weighted Gene Co-expression Network Analysis (WGCNA) (Materials and Methods; **Fig. 3E-F**). This approach clusters genes into modules based on correlated expression across samples, thereby revealing coordinated biological networks.

Hierarchical clustering identified several distinct co-expression modules (**Fig. 3E, Supplementary Table S3**), yet none showed statistically significant correlation with APAP exposure at any dose **(Fig. 3F**). These findings indicate that, while APAP exposure impacts the expression of a small number of genes, it does not disrupt broader transcriptional architecture or gene co-regulation patterns under physiologic exposure conditions in this CO system.

### Acetaminophen exposure does not alter progenitor or mature neuron proportions in organoids

Given the enrichment of neurogenesis-and neuronal differentiation-related pathways among APAP differentially expressed genes, we next examined whether APAP exposure altered progenitor proliferation or neuronal differentiation. To address this, we performed immunofluorescence staining on sections from 1-month-old organoids using Ki67 (proliferative progenitor marker), PAX6 (radial glia/neural progenitor marker), NeuN (post-mitotic neuronal marker), and DAPI (nuclei) (**Fig. 4A-C**).

**Figure 4.**
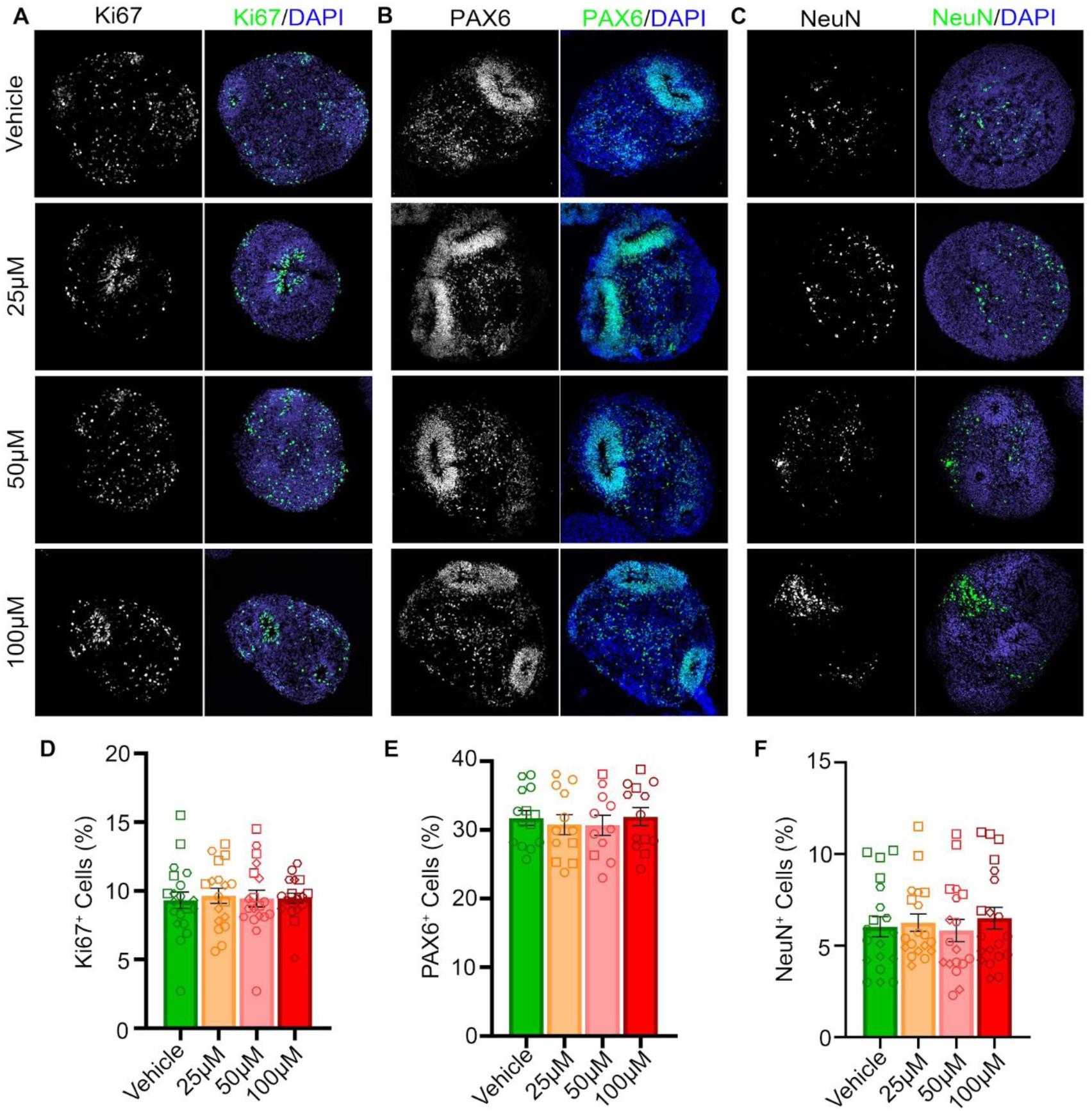
APAP exposure does not alter neural progenitor or mature neuron proportions in cortical organoids. **(A-C)** Representative confocal images of 20 μm cryosections from one-month-old organoids immunostained for Ki67 (green; **A**), Pax6 (green; **B**) or NeuN (green; **C**), with DAPI (blue) counterstain. Scale bar: 200μm. **(D-F)** Quantification of Ki67^+^ progenitor cells (**D**, n=4 iSPC lines, 3 independent batches), PAX6^+^ (**E**, n=3 iPSC lines, 3 independent batches) and NeuN^+^ neurons (**F**, n=4 iPSC lines, 3 independent batches) as a percentage of total cells across all exposure conditions. Data represent mean ± SEM (n≥10 organoids, n≥3 lines per marker, 3 independent batches per marker); data points represent individual organoids, different shapes represent individual cell lines. Statistical significance was assessed using a mixed-effects model (REML) with Dunnett’s multiple comparisons test versus vehicle control. No significant differences were detected across exposure conditions.

Quantitative analysis showed that Ki67⁺ proliferative progenitors comprised approximately 8-10% of total cells across all exposure conditions (**Fig. 4D**), while PAX6⁺ neural progenitors accounted for ∼30-40% of cells (**Fig. 4E**), with no detectable differences between conditions. NeuN⁺ neurons represented ∼5% of the total cell population (**Fig. 4F**). No significant differences were observed between vehicle-and APAP-treated organoids. Together with the preserved Nestin and MAP2 expression patterns (**Fig. 1E-F**), these results indicate that therapeutic APAP exposure does not measurably alter progenitor proliferation, progenitor abundance, or neuronal differentiation at this developmental stage in this CO system.

### Acetaminophen exposure does not alter cell type composition or transcriptional profiles in three-month-old organoids

To assess whether APAP exposure at day 21 affects long-term CO development, we performed single-nucleus RNA sequencing (snRNA-seq) of organoids at 3 months of differentiation. Sequencing quality was high across all samples and conditions (**Fig. S3**), with 55,665 nuclei retained after quality control and filtering (average 6,958 per sample). The analysis identified the expected spectrum of cell type populations across all APAP treatment conditions, consistent with normal organoid maturation (**Supplementary Table S4**). Importantly, cell-type proportions were computed on a per-sample basis (i.e., the fraction of nuclei belonging to each cell type within each biological replicate), ensuring that each sample contributed equally to downstream statistical testing and avoiding bias from unequal cell numbers across samples. In particular, we detected mature upper-layer excitatory neurons as well as precursors of inhibitory interneurons, reflecting the progressive emergence of cortical-like neuronal diversity (**Fig. 5A-B**). Quantification of cell-type proportions across exposure conditions revealed no significant differences, indicating that early APAP exposure did not significantly alter the relative composition of these populations at a later time point (**Fig. 5C**).

**Figure 5.**
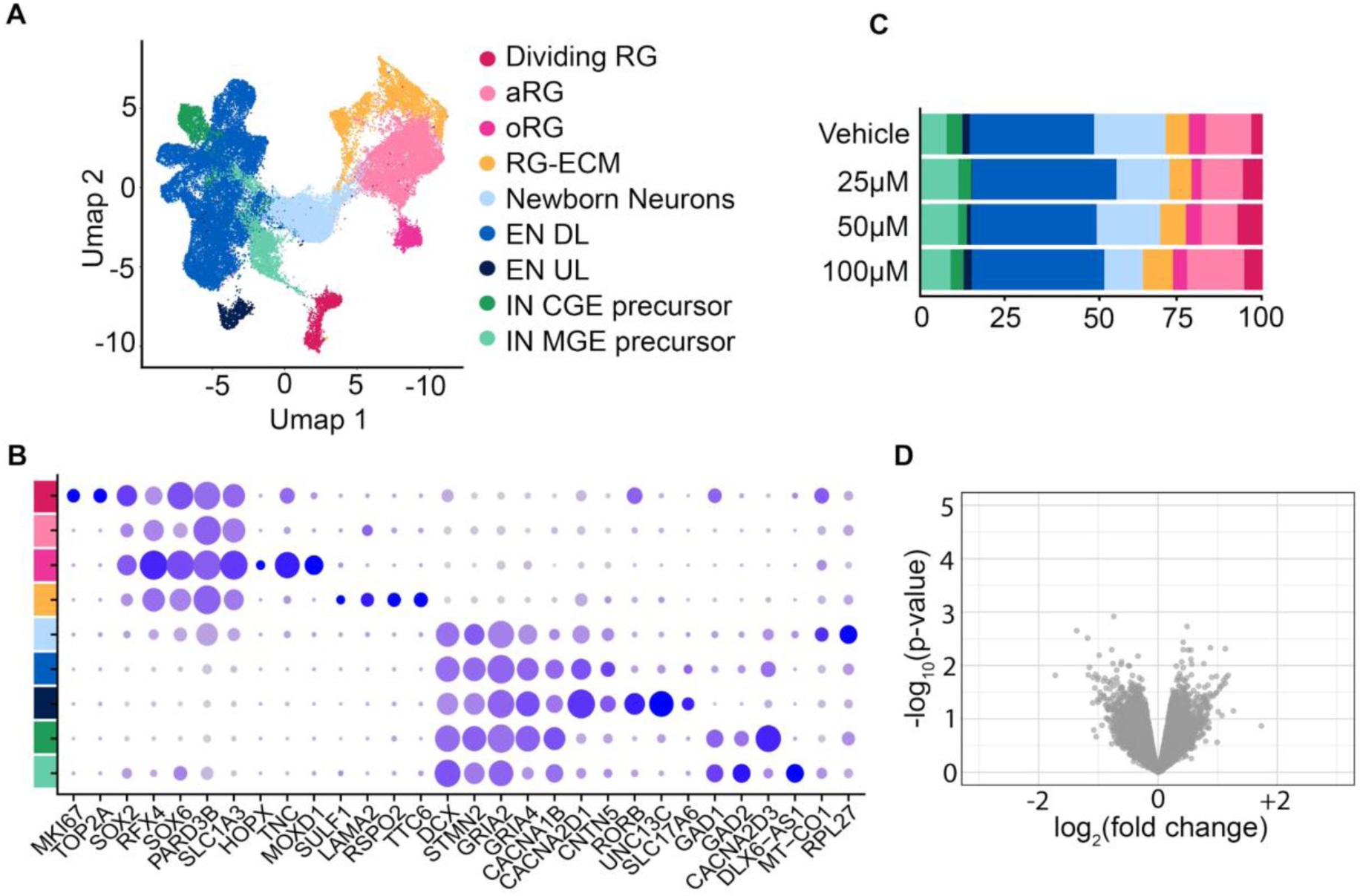
Single-nucleus RNA sequencing of 3-month-old organoids following short-term APAP exposure. **(A)** UMAP representation of snRNA-seq data showing major cell populations identified in organoids, including dividing radial glia (Dividing RG), apical radial glia (aRG), outer radial glia (oRG), radial glia–Extra cellular matric (RG-ECM), newborn neurons, upper-layer excitatory neurons (EN UL), deep-layer excitatory neurons (EN DL), caudal ganglionic eminence inhibitory precursors (IN CGE precursor), and medial ganglionic eminence inhibitory precursors (IN MGE precursor). **(B)** Dot plot of cell type markers. Dot size represents the proportion of cells expressing each gene, and color intensity represents average expression level. **(C)** Relative proportions of major cell populations across treatment conditions (Vehicle, 25 µM, 50 µM, 100 µM APAP), showing no significant differences in cell type composition (n=2 independent lines per condition). **(D)** Volcano plot of differential gene expression in representative EN DL cluster at 100 µM APAP exposure compared to vehicle. No genes reached statistical significance threshold (adjusted p-value < 0.05).

Pseudobulk differential gene expression analysis was performed both per cell-type cluster and across all cells combined. No DEGs were identified at any APAP dose or in any cell-type cluster, including EN DL, the largest and the best-powered cluster, at the highest concentration tested (100 µM) (**Fig. 5D, Supplementary Table S5**). Together, these results indicate that COs follow a typical trajectory of cellular differentiation and maturation, and that APAP exposure at day 21 does not induce persistent transcriptional or cellular changes detectable at 3 months within this CO model.

### Acetaminophen exposure does not alter electrophysiological activity or network connectivity in mature organoids

To determine whether APAP-induced transcriptional changes impacted neural function, we performed longitudinal electrophysiological recordings using MEAs on organoids from 2 months to 4 months of age. This method allowed us to evaluate spontaneous neural activity, network development, and potential long-term effects of APAP exposure.

We initially assessed whether COs were functionally coupled to the electrode arrays. All COs exhibited robust functional coupling to MEA electrodes across exposure conditions as indicated by consistent electrode resistance measurements (∼30 kΩ; **Fig. 6A-B**). By 4 months, all experimental groups displayed spontaneous electrophysiological activity with coordinated bursting patterns characteristic of mature neural networks (**Fig. 6C**).

**Figure 6.**
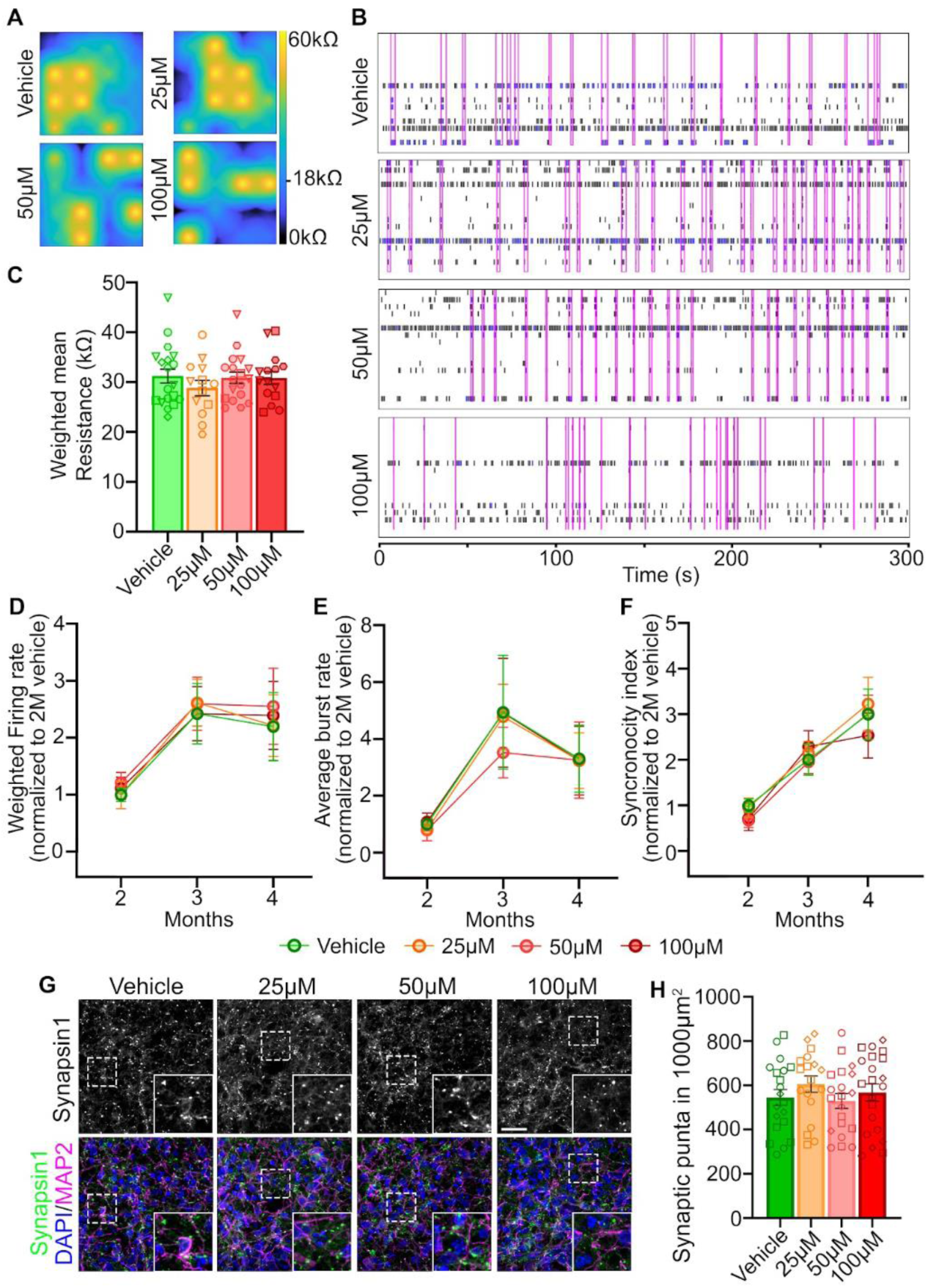
APAP exposure does not alter functional electrophysiological properties of COs. **(A)** Heat maps of electrode resistance measurements showing consistent organoid-electrode coupling across all exposure conditions. **(B)** Representative raster plots from four-month-old organoids showing spontaneous neural activity across conditions (vehicle, 25 μM, 50 μM, 100 μM APAP). Each horizontal trace represents activity from a single electrode, with black vertical tick marks indicating action potentials, and vertical magenta lines mark network activity. Time scale: 300 s. **(C)** Quantification of electrode resistance at 2 months demonstrates similar coupling across conditions. **(D-F)** Longitudinal analysis of firing rate **(D)**, burst frequency **(E)**, and synchronicity index **(F)** from 2 to 4 months. Values normalized to 2-month vehicle controls. Data represent mean ± SEM (n≥14 organoids, 5 iPSC lines, 3 independent batches). Statistical analysis: repeated-measures two-way ANOVA with Dunnett’s multiple comparisons versus vehicle control (all p > 0.05, not significant). **(G)** Representative confocal images of 20 μm cryosections from 3-months-old organoids immunostained for Synapsin1 (green), MAP2 (magenta) and DAPI (blue) counterstain. Scale bar: 20μm. **(H)** Quantification of synaptic puncta in a defined are of 1000um across all exposure conditions. Data represent mean ± SEM (n≥19 organoids, 3 iPSC lines, 3 independent batches); each data point represents an individual organoid; distinct symbols denote different cell lines. Statistical significance was assessed using a mixed-effects model (REML) with Dunnett’s multiple comparisons test versus vehicle control. No significant differences were detected across exposure conditions

Quantitative analysis revealed comparable electrophysiological maturation across exposure conditions. Firing rates increased from 2 to 4 months, as expected, with no differences between groups (**Fig. 6D**). Burst frequency analysis showed similar patterns of local neural activity, characterized by periods of high-frequency firing and burst generation, across time points and exposure conditions (**Fig. 6E**). Synchrony index values, reflecting coordinated network activity across electrodes, likewise remained unchanged across doses and timepoints (**Fig. 6F**).

To further assess synaptic maturation, we performed immunostaining for the synaptic marker Synapsin1 in 3-month-old organoids (**Fig. 6G**). Quantification revealed no differences in synaptic puncta density across exposure conditions (**Fig. 6H**). Together, these results indicate that APAP exposure does not measurably affect spontaneous firing, local bursting, or network-level coordination, along with synaptic puncta density, in cortical organoids and are in line with prior studies in 2D and 3D human neuronal culture models in which APAP was used as a negative control and did not affect baseline electrophysiological activity^24–27^

## DISCUSSION

In this study, we used human induced pluripotent stem cell-derived cortical organoids to examine the effects of acetaminophen exposure on early cortical development *in vitro*. Across physiologically relevant concentrations, APAP exposure did not significantly alter organoid growth, cytoarchitecture, neuronal differentiation, cell type composition, or network activity. Only modest transcriptional changes were observed by bulk but not by the single cell RNAseq. Furthermore, these molecular changes were not accompanied by detectable alterations in progenitor or neuronal populations, synaptic maturation, or electrophysiological network properties.

The preservation of cellular and functional outcomes despite modest gene expression shifts is consistent with the plasticity and homeostatic compensation of developing neural circuits^48^. Thus, our findings indicate that, within the exposure window and experimental system tested here, therapeutic-range APAP exposure does not produce detectable large-magnitude effects on early human cortical developmental processes.

An important consideration is that cortical organoids primarily model direct exposure of developing cortical tissue to the parent APAP compound, rather than systemic APAP metabolism. APAP metabolite-mediated effects require enzymatic pathways that are not strongly represented in our model, consistent with the limited expression of these pathways in early fetal cortex. AM404 formation requires FAAH-dependent metabolism, and FAAH transcript expression was low in our organoids, consistent with BrainSpan^29,49^ fetal brain samples from the corresponding developmental periods (**Fig. S4**). Likewise, NAPQI formation requires CYP-mediated oxidation of APAP, yet the relevant CYP enzymes, including CYP2E1, CYP1A2, and CYP3A4, were low or undetectable in both organoids and fetal brain samples (**Fig. S4**). Expression of APAP conjugation and detoxification genes, including UGT, SULT, and GST family members, was also broadly similar between organoids and fetal brain (**Fig. S4**). Consistent with these data, pregnancy PBPK modeling predicts very limited fetal conversion of APAP to NAPQI, with a median molar dose fraction of only 0.06% in the term fetus^45^. Together, these findings suggest that cortical organoids resemble early fetal brain in having limited predicted capacity for local APAP metabolism to AM404 or NAPQI. Thus, our data address direct effects of parent APAP exposure under the conditions tested, but do not exclude metabolite-mediated and systemic effects that could arise *in vivo* through maternal, placental, or hepatic metabolism.

Our findings differ from some rodent studies reporting APAP-induced apoptosis and behavioral effects^20,21^, highlighting known differences between experimental systems, developmental timing, and species-specific drug metabolism^23^. In rodents, APAP-associated neurotoxicity has been attributed in part to formation of the reactive metabolite N-acetyl-p-benzoquinone imine (NAPQI), generated through cytochrome P450-mediated oxidation ^50^. In humans, fetal CYP2E1 expression is minimal until late gestation and increases primarily after birth^51,52^ with APAP detoxification during early development occurring predominantly via sulfation pathways^53^. Because cortical organoids lack hepatic metabolism, the responses observed here likely reflect direct effects of the parent APAP compound within this *in vitro* system, rather than metabolite-mediated toxicity.

Our findings also provide experimental context for recent sibling-controlled epidemiological studies suggesting that previously reported associations between prenatal APAP exposure and neurodevelopmental outcomes may be influenced by confounding factors^17–19^. However, our study does not address clinical neurodevelopmental outcomes directly. Rather, it provides a controlled in vitro test of whether therapeutic-range APAP exposure produces detectable effects on early human cortical developmental phenotypes.

Our study has several limitations. First, APAP exposures were short-term and designed to approximate typical maternal use; the effects of prolonged or supratherapeutic dosing were not examined. Second, although effect sizes across assays were consistently small and not statistically significant, we cannot exclude subtle biological effects below the sensitivity of the current experimental system. We therefore interpret our findings as indicating a lack of detectable or large-magnitude effects under the conditions tested, rather than definitive absence of any effect. Third, cortical organoids lack vascular, immune, placental, and peripheral metabolic components that may influence APAP pharmacokinetics and pharmacodynamics *in vivo*; future studies could incorporate microfluidic platforms or co-culture systems with vascular, glial, immune, placental, or hepatic components to address these factors. Fourth, functional assessment was limited to multielectrode array recordings up to 4 months, and additional approaches such as calcium imaging, proteomics, or extended network-level assays may reveal more subtle phenotypes. Finally, we did not evaluate combined exposures, such as APAP with other medications or environmental factors, nor did we assess responses in organoids carrying ASD-associated genetic variants; therefore, potential gene-environment interactions or genotype-specific effects were not examined in this study.

Despite these caveats, our findings add to a growing body of experimental work examining APAP exposure in human-relevant *in vitro* models and indicate that, within this cortical organoid system, therapeutic APAP exposure does not measurably perturb early cortical developmental processes. Future studies should extend exposure paradigms, incorporate metabolite-specific assays and neuroimmune models, and evaluate gene-environment interactions using organoids carrying neurodevelopmental risk variants.

In conclusion, therapeutic APAP exposure in this cortical organoid model produced only minor transcriptional changes without detectable effects on cellular composition, structural maturation, or network function. Together with prior studies in human and animal experimental systems^24–27^, these findings place our results within ongoing efforts to use controlled, human-relevant models to interpret epidemiological observations, while avoiding conclusions beyond the scope of this *in vitro* system.

## SUPPLEMENTARY FIGURES

**Supplementary Figure S1.**
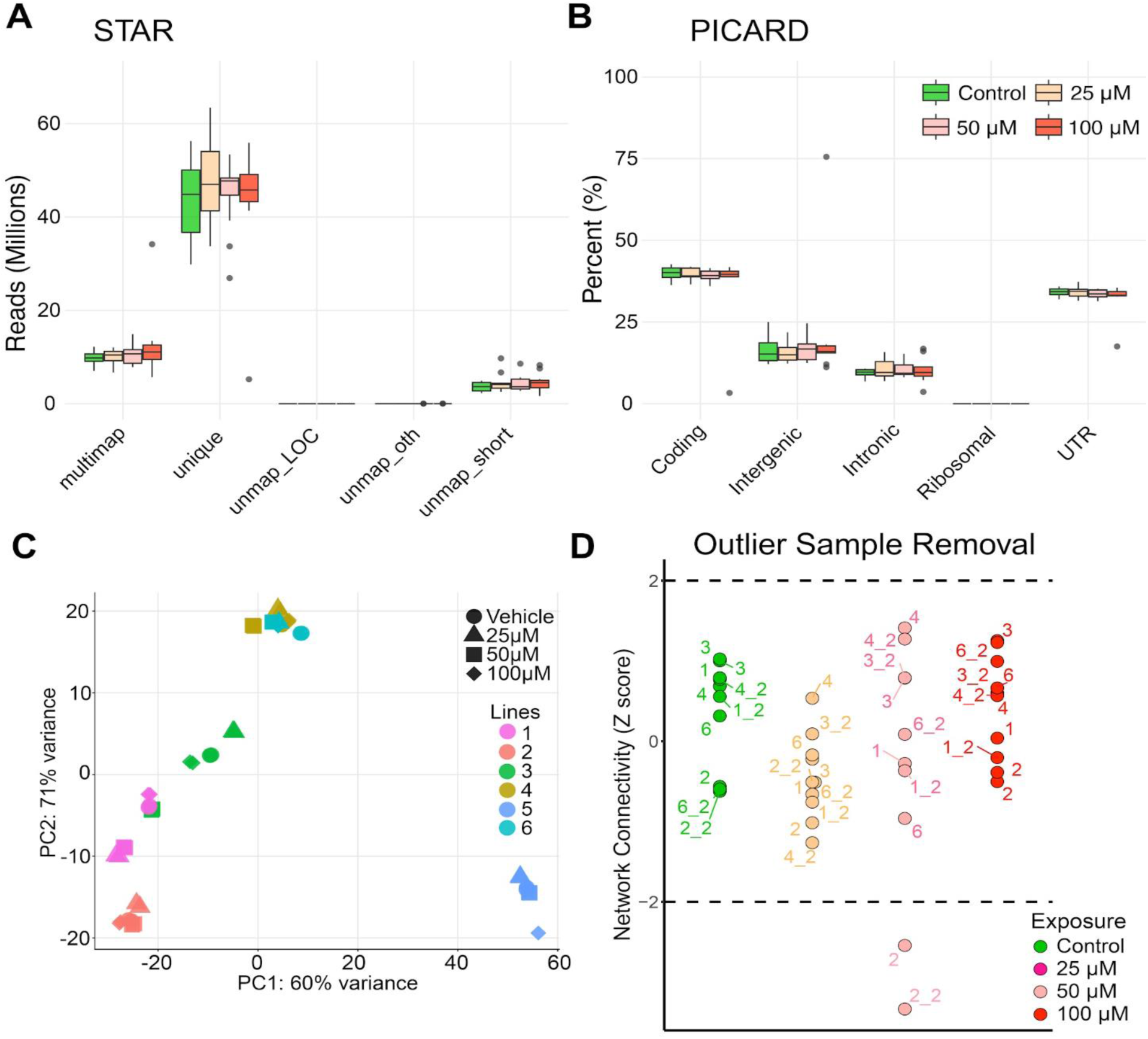
Quality control metrics for RNA-seq data. (**A**) Sequencing metrics from STAR (v2.5.3a) for each condition at 1 month (Vehicle, 25μM, 50μM, and 100μM). (**B**) Sequencing metrics from PicardTools (v2.12) for each condition at 1 month. (**C**) PCA of organoid samples derived from 6 iPSC lines. Line 5 was removed from the subsequent experiments as an outlier. **(D)** Sample outlier removal was performed with the WGCNA package in R for each condition at 1 month based on Z-scores of standardized network connectivity. Outliers were defined as samples with Z scores of <|-2|.

**Supplementary Figure S2.**
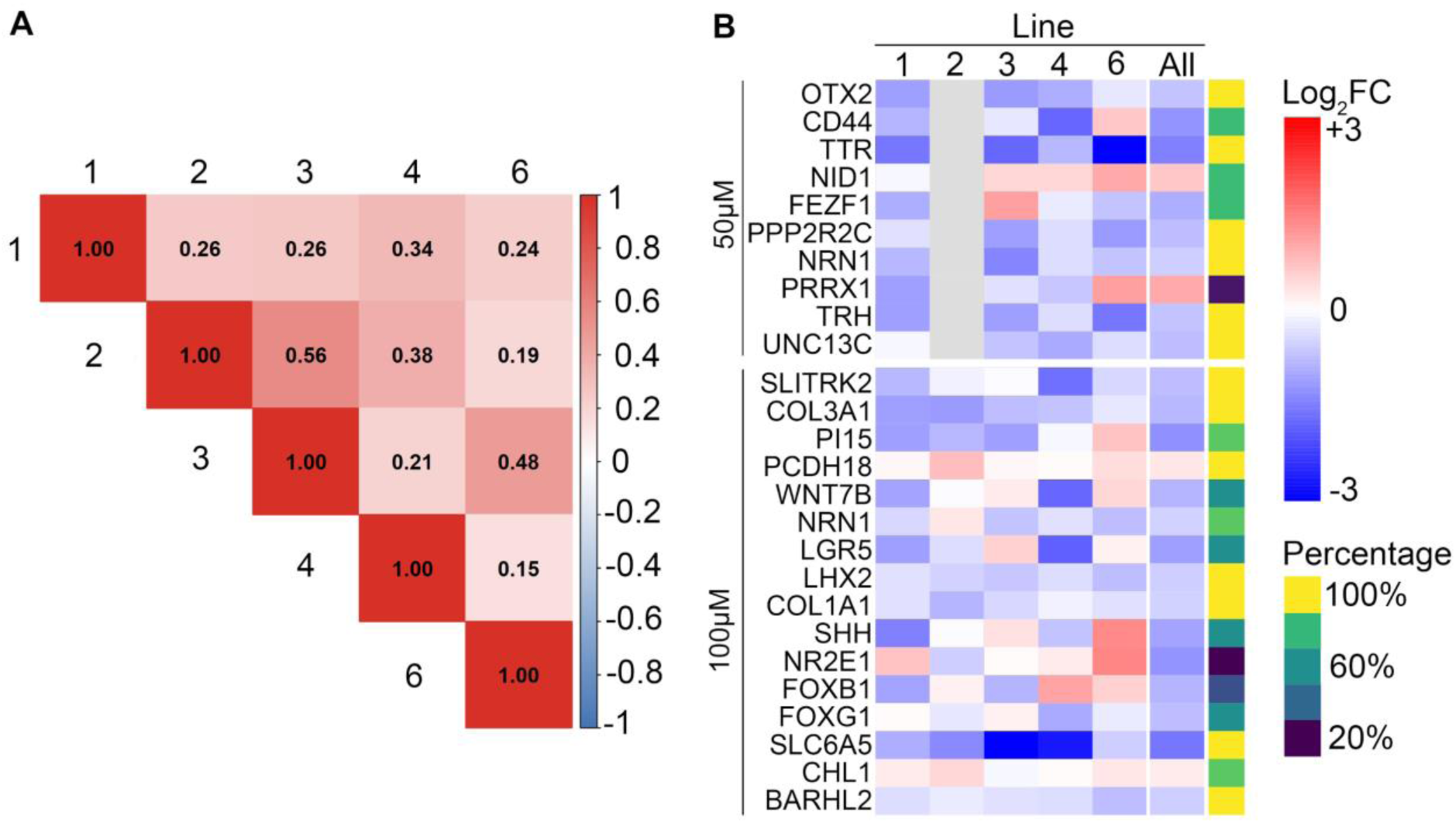
Cross-line correlation of differentially expressed genes identified by bulk RNA-seq. **(A)** Pearson correlation matrix showing the concordance of DEG log2 fold-change values across individual hiPSC lines. Correlations were calculated using the log2 fold-change (FC) values of genes identified as differentially expressed. Values indicate Pearson correlation coefficients (r) between lines. **(B)** Heatmap displays log2 fold change (APAP vs. control) for each DEG (adjusted p < 0.05, |log2FC| > 0.2) across individual hiPSC lines (1, 2, 3, 4, 6) and the pooled analysis (All), separated by dose (50 µM, top; 100 µM, bottom). Color scale ranges from −3 (blue, downregulated) to +3 (red, upregulated). Gray cells indicate missing data due to QC-based outlier exclusion (line 2 at 50 µM; see **Fig. S1**). The sidebar indicates the proportion of lines with fold changes in the same direction as the pooled result (yellow = 100%, teal = 60%, purple = 20%).

**Supplementary Figure S3.**
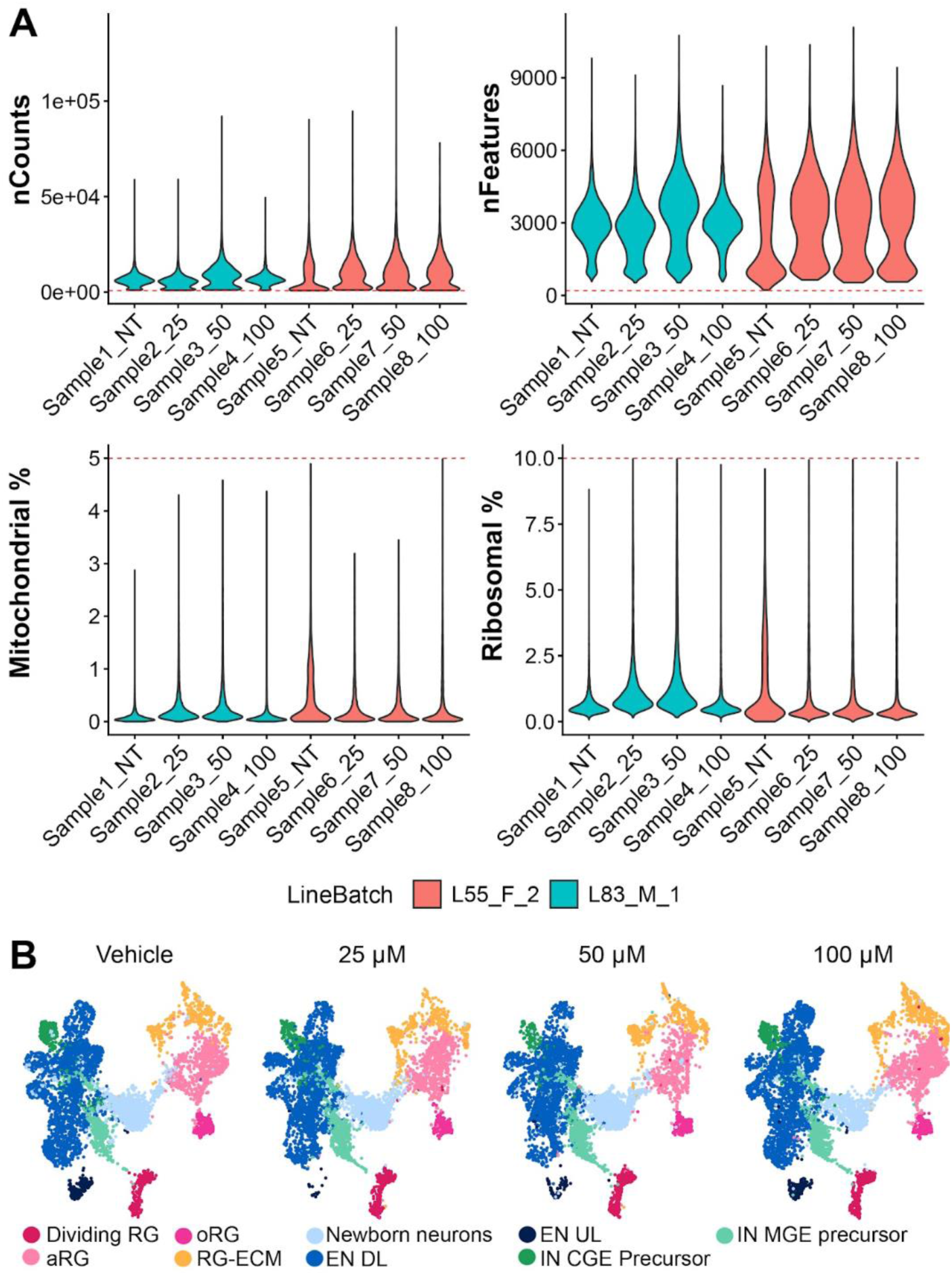
Quality control metrics and integration of snRNA-seq data from 3-month-old COs. **(A)** Violin plots showing per-sample distribution of key quality control metrics: unique molecular identifiers (nCounts), genes detected per nucleus (nFeatures), mitochondrial gene content (Mitochondrial %), and ribosomal gene content (Ribosomal %). Dashed red lines indicate filtering thresholds (nCounts > 800, nFeatures > 200, mitochondrial < 5%, ribosomal < 10%). Samples are colored by cell line and batch (L55_F_2, salmon; L83_M_1, teal). **(B)** UMAP projections of Harmony-integrated snRNA-seq data split by APAP treatment condition (Vehicle, 25 µM, 50 µM, 100 µM), colored by annotated cell type. Cell populations include dividing radial glia (DivRG), apical radial glia (aRG), outer radial glia (oRG), radial glia–extracellular matrix (RG-ECM), newborn neurons, upper-layer excitatory neurons (EN UL), deep-layer excitatory neurons (EN DL), mature deep-layer excitatory neurons (Mature EN DL), LHX9-expressing deep-layer excitatory neurons (EN DL LHX9), caudal ganglionic eminence inhibitory precursors (IN CGE), and medial ganglionic eminence inhibitory precursors (IN MGE). UMAPs are shown on equally down-sampled nuclei across samples.

**Supplementary Figure S4.**
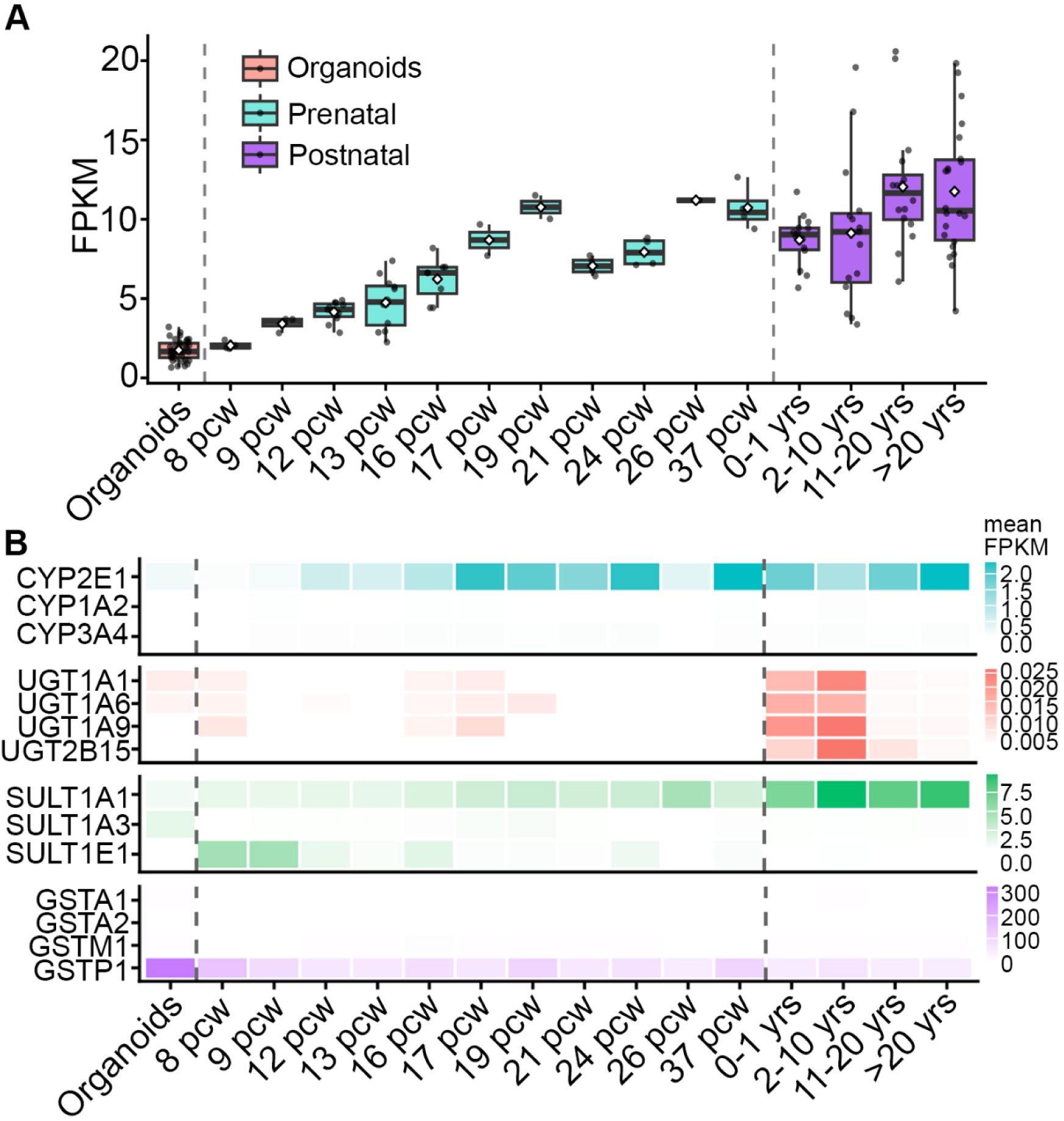
Expression of genes implicated in APAP metabolism across cortical development. **(A)** FAAH FPKM in 1month old cortical organoids (salmon) and human cortex from BrainSpan at fetal (pcw, turquoise) and postnatal (purple) timepoints. Diamonds indicate the mean. Dashed lines separate organoids from fetal brain and fetal from postnatal samples. pcw, post-conceptional weeks. **(B)** Heatmaps show mean FPKM of CYP450 (oxidative metabolism), UGT (glucuronidation), SULT (sulfation), and GST (glutathione conjugation) gene families in 1 month old cortical organoids and human cortex (averaged across 6 cortical regions) from BrainSpan across fetal (pcw) and postnatal timepoints. Each gene family is shown with a different color scheme. Dashed lines separate organoids from fetal brain and fetal from postnatal samples. pcw, post-conceptional weeks.

## METHODS

### iPSCs lines

hiPSC lines used in this project (**Supplementary Table S6**) were derived from typical developing “control” individuals as previously described^37,38,54,55^. Before thawing, tissue culture dishes were coated with Matrigel (1:200), diluted in Dulbecco’s modified Eagle medium (DMEM)/F12 (GiCO), and left at 37°C for at least 2 h. Frozen hiPSCs, stored in CryoStor (BioLife Solutions), were quickly thawed in a water bath at 37°C for 45 s and immediately rinsed with prewarmed mTeSR Plus medium (StemCell Technologies). Cells were centrifuged (3 min, 200 x g), and the pellet was resuspended in 3mL mTeSR plus supplemented with 5μM ROCK Inhibitor (RI; Y-27632 StemCell Technologies, cat. 72304). Cells were plated on Matrigel-coated plates and maintained at 37°C, 5% CO_2_. After 3 h, the medium was replaced with 3mL of fresh mTeSR Plus media (without RI). Medium was subsequently changed every other day. For routine passaging, iPSC colonies were manually selected and transferred to new Matrigel-coated plates every 4 to 6 days. Briefly, the culture medium was aspirated, and the plate was rinsed with 1 mL of fresh mTeSR Plus. Using a sterile 10 μL pipette tip, ∼20 undifferentiated colonies were carefully scraped and dissociated by gentle pipetting in 2 mL of fresh prewarmed mTeSR Plus medium. Finally, the resulting suspension was transferred to a new Matrigel-coated plate. To maximize recovery, an additional 1 mL of mTeSR Plus medium was used to rinse the original plate and collect any remaining cells. Cultures were then maintained at 37°C, 5% CO_2_.

### Generation of cortical organoids

Cortical organoids were generated from hiPSCs using the semi-guided protocol described^36,37^. hiPSCs (at approximately 80% confluency) were exposed to 10μM RI ∼1 hr before beginning organoid preparation. Subsequently, cells were dissociated using a 1:1 solution of Accutase (MP Biomedicals) and Phosphate-buffered Saline (PBS; Corning), and incubated at 37°C with gentle mechanical mixing every 10 to up to 50 min. The cell suspension was diluted in PBS and passed through a 40 μm cell strainer (Corning) to obtain a single-cell suspension, followed by centrifugation (3 min, 200 x g). The cell pellet was resuspended in mTeSR Plus medium supplemented with 5 μM ROCK inhibitor, 1 μM Dorsomorphin (Dorso; Tocris), and 10 μM SB431542 (SB; Stemolecule). A total of 4x10^6^ cells were seeded per well of a 6-well plate (Genclone) in 3 mL of this medium for 24 h under rotation (95 rpm) at 37°C, 5% CO_2_ to promote the formation of free-floating spheres.

Medium was changed daily and replaced with mTeSR Plus supplemented with 1 μM Dorso (Tocris) and 10 μM SB (Stemolecule) until day 4. From days 4 - 8, medium was changed every other day with Media 1 (Neurobasal medium (GiCO), 1x GlutaMAX (GiCO), 1% MEM non-essential amino acids (NEAA; GiCO), 1% Penicillin-Streptomycin (PS; GiCO), 2% Gem21 (Gemini), and 1% N2 Neuroplex (Gemini) supplemented with 1 μM Dorso and 10 μM SB. From days 9 to 15, the medium was changed daily and replaced with Media 2 (Neurobasal medium,1x GlutaMAX, 1% NEAA, 1% Pen-Strep, and 2% Gem21) and 20 ng/mL fibroblast growth factor-2 (FGF2; R&D Systems). From days 16 - 21, the media was changed every other day with Media 2 supplemented with 20 ng/mL FGF2 and 20 ng/mL epidermal growth factor (EGF; Peprotech). From days 22 - 27, medium was changed every other day with Media 2 supplemented with 10 ng/mL each of brain-derived neurotrophic factor (BDNF; Peprotech), glial cell-derived neurotrophic factor (GDNF; Peprotech), and neurotrophin-3 (NT3; Peprotech), 200 μM L-Ascorbic Acid (Sigma-Aldrich), and 1 mM dibutyryl-cAMP (dCAMP; StemCell Technologies). From days 28 - 30, the medium was changed every other day with Media 2. For long-term culture, organoids were maintained in Media 2 with medium changes every 3-4 days. Unusually small or unhealthy (dark) organoids were excluded from the subsequent analyses, as they do not represent properly developed organoids.

### Acetaminophen Exposure

APAP exposure was initiated on Day 21 of the COs protocol. Stock solutions were prepared by diluting APAP in PBS. In 6-well plates, four wells were randomly assigned to one of four conditions: Vehicle (PBS), 25 μM, 50 μM, and 100 μM APAP. To achieve the desired concentrations, 3 μL of the corresponding APAP stock solution or PBS control was added to each well containing 3 mL of culture medium. APAP exposure was administered at the time of medium changes for 5 days, until day 26.

### Organoid Size Analysis

To assess the impact of APAP exposure on organoid growth, bright-field images of organoids were taken on days 21, 23, 25, 27, 29, and 31, as well as at 3 months, using an inverted microscope (EVOS FL Digital) with a 1.25x objective. Organoid size was measured using the Fiji-ImageJ software. Briefly, images were adjusted for contrast, and a threshold was applied to create a binary mask. The ‘Analyze Particles’ plugin was used to measure organoid area, applying a size filter of ∼20,000-infinity μm² and circularity of 0.7-1.0 to exclude debris. Low-quality organoids were manually removed from analyses. For growth evaluation, average organoid size was calculated for each condition at each time point and normalized to the measurement taken on day 21.

### Organoid Dissociation and Apoptosis Assay

To establish a positive control, a subset of unexposed organoids was treated with 1 μM camptothecin (CPT; Thermo Scientific), a well-characterized inducer of apoptosis, 18 h before the assay. On day 26, CPT-exposed organoids and APAP-exposed organoids were harvested and dissociated into single cells.

COs (10–12 per sample) were collected and dissociated into individual cells for the apoptosis assay. After two rinses with PBS containing 0.1% bovine serum albumin (BSA; Fisher Scientific), the organoids were incubated at 37°C for up to 50 mins in a dissociation solution containing Accutase, papain (16.7 U/ml; Worthington), and Deoxyribonuclease I (100 U/ml; Worthington) in Hank’s Balanced Salt Solution (HBSS; ThermoFisher).

To assist with dissociation, the cell suspension was gently pipetted every 10 minutes. Once dissociation was achieved, warm M2 medium was added to stop enzymatic activity. The suspension was then filtered through a 40 μm cell strainer, pre-wetted with 0.1% BSA/PBS solution, centrifuged for 7 mins at 200 x g, and the resulting cell pellet was resuspended in pre-chilled PBS containing 0.1% BSA.

A single-cell suspension from each condition was quantified using a Countess 3 Automated Cell Counter (Thermo Fisher) and diluted to 3 × 10^5^ cells/mL. Triplicate wells of a 96-well plate (Greiner) were seeded with 3 × 10⁴ cells per well. Each well received an equal volume of Caspase-Glo® 3/7 3D Reagent (Promega), resulting in a 1:1 ratio of cell suspension to reagent.

Plates were processed on a Spark Multimode Microplate Reader (Tecan) using the following protocol: (1) orbital shaking at 510 rpm for 30 s to ensure mixing; (2) 30 min incubation to allow caspase-mediated cleavage of the luminogenic substrate; and (3) luminescence measurement. The luminescence signal was proportional to caspase-3/7 activity, providing a quantitative readout of apoptosis.

### RNA extraction and sequencing

Total RNA was extracted from 1-month old COs from a total of 48 samples (4 conditions: vehicle, 25, 50, and 100 µM APAP × 6 lines × 2 technical replicate per condition, with replicates generated from independent organoid batches grown on different days) using the RNeasy Kit (QIAGEN). Ribosomal RNA was depleted with the Ribo-Zero rRNA Removal Kit (Illumina), and libraries were prepared using the TruSeq Stranded Total RNA Kit. Paired-end (100 bp) sequencing was performed on an Illumina NovaSeq S4 platform to a mean depth of ∼25 million reads per sample.

### Data processing and normalization

Reads were trimmed for adapters and low-quality bases (Cutadapt v1.18) and aligned to the human genome (GRCh38.p13, Gencode v42) using STAR (v2.7.10b). Gene-level quantifications were generated with RSEM (v1.2.28). QC metrics were summarized with RNA-SeQC, featureCounts, PicardTools, and Samtools. Expressed genes were defined as genes with TPM > 0.2 in at least 25% of samples within each exposure group (Vehicle, 25µm, 50µm, 100µm). Outliers (Z < –2) were removed, and surrogate variables were estimated with SVA. VariancePartition and MARS analyses informed final covariate selection.

### Differential gene expression and weighted gene co-expression network analysis

Differential expression was assessed using limma-voom with duplicateCorrelation, including exposure, cell line, and selected surrogate variables. Genes meeting the differential expression criteria (log2 fold change ≥ 0.2 and FDR-adjusted p < 0.05) were considered as differentially expressed. Gene annotations were retrieved via biomaRt. Weighted gene co-expression network analysis (WGCNA) identified modules of co-expressed genes using blockwiseModules function with the following parameters: corType=cor; networkType=signed; pamStage = TRUE; reassignThreshold = 1e-6; mergeCutHeight = 0.25; maxBlockSize = 16000; Soft threshold power = 10; deepSplit = 0; minModuleSize = 150. Module eigengene-exposure correlations were calculated with linear models including cell line as a covariate. Multiple-testing correction across all modules was performed using the Benjamini–Hochberg FDR procedure, and modules with FDR < 0.05 were considered significantly associated with APAP exposure.

### Nuclei isolation and snRNAseq library preparation

Single-nucleus RNA sequencing (snRNA-seq) was performed on two hiPSC-derived organoid lines using the PIP platform (Illumina) following the manufacturer’s instructions. Nuclei were isolated from 3-month-old cortical organoids (a total of 8 samples: 2 lines x 4 conditions: vehicle, 25, 50, and 100 µM APAP) using a mild detergent-based lysis buffer consisting of PBS, 10 mM NaCl, 3 mM MgCl₂, 1 mM DTT, 0.1% Tween-20, 0.1% IGEPAL CA-630, 0.01% BSA, and 1 U/µL RNase inhibitor. Organoids were incubated in a lysis buffer for 5–10 minutes to allow gentle membrane disruption. Lysis was stopped by adding an equal volume of wash buffer (PBS, 10 mM Tris-HCl pH 7.4, 3 mM MgCl₂, 0.01% BSA, 1 U/µL RNase inhibitor), followed by sequential filtration through 75-µm and 40-µm strainers to remove debris. The nuclei suspension was centrifuged at 500 × g for 5 minutes at 4 °C and washed twice with wash buffer under the same conditions. The final pellet was gently resuspended in nuclei resuspension buffer (PBS, 0.1% BSA, 1 U/µL RNase inhibitor). Nuclei concentration and integrity were assessed using ethidium homodimer.

Library preparation was performed using the Illumina Single Cell 3′ RNA Prep workflow with Particle-Templated Instant Partitions (PIPseq™ chemistry). Briefly, cells were combined with barcoded hydrogel particles and emulsified to capture mRNA. After lysis and reverse transcription, cDNA was amplified and processed into sequencing libraries, which were sequenced on an Illumina NovaSeq S4 platform.

### Single-nucleus RNA-seq data processing and analysis

Raw FASTQ files were processed using the DRAGEN™ Single Cell pipeline to generate gene-by-cell count matrices for downstream analysis using the Seurat package (v5) in R. Low-quality nuclei were removed based on the following thresholds: nCount < 800, nFeature < 200, mitochondrial fraction > 5%, and ribosomal fraction > 10%. Doublets were identified and removed using doubletFinder. Each sample was normalized using SCTransform, and the merged dataset was subjected to principal component analysis, followed by batch correction with Harmony using the LineBatch variable to account for line-specific variation. Dimensionality reduction was done using RunUMAP, and graph-based clustering was performed using FindNeighbors and FindClusters with a resolution of 0.2. Cluster differential gene expression analysis was performed using FindAllMarkers and cell-type annotation was based on canonical marker genes.

Changes in cell type proportions were assessed using Propeller, which applies a logit transformation followed by linear modeling to account for compositional data. Importantly, cell-type proportions were computed on a per-sample basis (i.e., the fraction of nuclei belonging to each cell type within each biological replicate), ensuring that each sample contributed equally to downstream statistical testing and avoiding bias from unequal cell numbers across samples. Raw numbers of nuclei per sample and per cluster are reported in **Supplementary Table S4**. Three complementary approaches were used: (1) Pairwise comparisons testing each APAP dose against the non-treated control using robust t-tests with propeller.ttest, (2) Dose-response trend analysis testing for linear dose-dependent changes by modeling dose as a continuous variable (0, 25, 50, 100) using limma’s lmFit and eBayes functions with robust variance estimation, and (3) Overall dose effect testing whether any APAP dose differed from control using an F-test across all three pairwise contrasts simultaneously via contrasts.fit. Differential gene expression analysis was performed using the Dreamlet pipeline, which aggregates counts to pseudobulk by sample and cell type, partitions variance to identify key covariates (LineBatch and nCounts), and models the effects of APAP exposure. Contrasts included each dose versus control and pairwise comparisons between doses.

### Immunofluorescence staining and imaging

Cortical organoids were collected, rinsed briefly with PBS, and fixed in 4% paraformaldehyde (PFA) overnight at 4°C. After fixation, organoids were washed three times in PBS and transferred to a 30% sucrose solution for cryoprotection. The organoids were kept at 4°C until they sank to the bottom of the tubes, indicating complete sucrose infiltration. They were then transferred to a cryomold (Fischer Scientific), embedded in Tissue-Tek OCT medium (Sakura), and frozen on dry ice. Cryosections (20μm thick) were prepared using Leica CM1950 cryostat at -20°C and mounted on SuperFrost Plus slides (Fisher Scientific).

For immunostaining, slides were washed three times with PBS (10 mins each) to remove residual OCT. Permeabilization and blocking were performed simultaneously by incubating slides for at least 1 h at room temperature in blocking solution [5% Horse serum (HS, GiCO) and 0.1% Triton X-100 (Sigma Aldrich) in PBS]. Following blocking, slides were incubated overnight at 4°C with primary antibodies diluted in the same blocking solution.

The following primary antibodies were used: SOX2 (mouse, 1:200, Abcam), NeuN (rabbit, 1:200, Cell Signaling), MAP2 (chicken, 1:3000, Abcam), Ki67 (rabbit, 1:500, Abcam), Nestin (mouse, 1:200, Abcam), ZO1 (mouse, 1:500, BD Biosciences), and Cleaved Caspase-3 (rabbit, 1: 400, Cell Signalling).

After overnight incubation with primary antibodies, slides were washed three times with PBS (10 mins each) to remove unbound antibodies. Sections were then incubated for 1 hr at room temperature with species-specific Alexa Fluor-conjugated secondary antibodies (1:1000 dilution in blocking solution). The following secondary antibodies were used: anti-rabbit Alexa Fluor (488), anti-mouse Alexa Fluor (555), and anti-chicken Alexa Fluor (647) (Abcam). Following secondary antibody incubation, sections were washed three times with PBS (10 mins each) and counterstained with DAPI (1μg/1mL, Thermo Scientific) for 10 mins at room temperature. Slides were washed three additional times with PBS (10 mins each) before being mounted using ProLong Gold antifade reagent (Life Technologies).

Immunostained CO sections were imaged using a Nikon A1 confocal microscope with appropriate laser lines and filter sets. For each condition, at least four organoids were analyzed. Image analysis was performed using Fiji-ImageJ, an open-source software based on ImageJ. Due to the dense and complex architecture of organoid slices, nuclear segmentation was not feasible; therefore, nuclear counting was avoided. Instead, marker colocalization with DAPI-stained nuclei was quantified using the ‘Just Another Colocalization Plugin’ (JaCoP) in ImageJ. Manders’ coefficient was used to assess the degree of overlap between marker and nuclear signals, providing a robust measure of marker-positive cells within highly packed tissues.

### Multielectrode Array (MEA) Recording

At day 45, COs were plated onto 48-well multielectrode array (MEA) plates (Axion Biosystems). MEA plates were coated using a two-step process before plating. First, 100 μL of poly-L-ornithine hydrobromide (10 μg/mL in PBS; Sigma-Aldrich) was added to each well and incubated overnight at 37°C. Wells were then rinsed three times with sterile MilliQ water containing 1% PS. Next, 100 μL of Laminin (Engelbreth-Holm-Swarm murine sarcoma basement membrane; Sigma-Aldrich) diluted 1:300 in PBS was applied to each well and incubated overnight at 37°C.

Following coating, 2-3 organoids were carefully transferred into each well using a wide-bore pipette tip and positioned over electrodes under a microscope. Each well was initially filled with 250 μL of Media 2. With the next two medium changes, an additional 250 μL of fresh Media 2 was added per well, until a final volume of 750 μL was reached. For all subsequent medium changes, one-third (250 μL) of the culture volume (250 μL) was slowly aspirated and replaced with an equal volume of fresh Media 2. Medium was changed twice a week (every Monday and Friday).

At day 60 of culture, the medium change regimen (replacing one-third of the volume at each change) was maintained, but Media 2 was gradually phased out and replaced with BrainPhys Neuronal Medium (StemCell Technologies) supplemented with 1% NEAA, 1% PS, 2% SM1 (StemCell Technologies), and 1× GlutaMAX.

Weekly electrophysiological recordings were performed using the Maestro Pro MEA system (Axion Biosystems) at approximately the same time each week. Before recording, MEA plates were acclimated to 37°C in the Maestro device for 5 mins, followed by a 5 min recording period. Spontaneous activity was sampled at a rate of 12.5 kHz and filtered using a Butterworth band-pass filter (200 Hz - 3 kHz). Spikes were detected using an adaptive threshold set at 5.5 x the noise standard deviation. Data was analyzed with Axion’s AxIS Navigator software (version 2.5.2; Axion Biosystems). Electrodes were classified as active if they exhibited ≥ 5 spikes/min. Bursts were defined as having an inter-spike interval threshold of ≥ 5 spikes within 100 ms, and network activity was considered present when at least 25% of active electrodes displayed bursting.

### Statistical Analysis and Visualization

All statistical tests and analyses are described in figure legends. GraphPad Prism (version 10.1.1) was used for statistical analysis and visualization of morphological, growth and cellular composition data (**Figs. 1, 2, 4, 6**). For datasets containing multiple organoids derived from the same iPSC line, statistical comparisons were performed using mixed-effects models (REML) with iPSC line included as a random effect to account for biological variability between independent cell lines. The complete experimental design, including hiPSC lines, differentiation batches, and assays performed for each experiment, is provided in the **Supplementary Table S7**. Source data and statistical analyses underlying main figures are provided in the **Supplementary Table S8**.

For bulk differential gene expression, genes with log2 fold change ≥ 0.2 and FDR-adjusted p < 0.05 considered as differentially expressed. For snRNAseq, differential gene expression analysis was performed using the Dreamlet with key covariates (LineBatch and nCounts). Pseudobulk differential gene expression analysis was performed both per cell-type cluster and across all cells combined. Gene Ontology enrichment analysis was conducted using g:Profiler with the human database as a background.

## ACKNOWLEDGEMENTS

The authors acknowledge funding support from the National Institutes of Health to L.M.I and/or A.R.M: 1R21HD109616, R56MH128365, R01ES033636, R21MH128827, 1OT2OD040415, and from California Institute for Regenerative Medicine CIRM DISC4-16377. We thank Dr. Zeyan Liew, as well as all members of Iakoucheva and Muotri labs for helpful discussions.

## DATA AVAILABILITY

Bulk and snRANseq data were deposited to GEO with the accession number GSE335987. The code is available at IakouchevaLab GitHub.

## Supplementary Table Titles

**Supplementary Table S1.** Differential gene expression (DEG) results.

**Supplementary Table S2.** Gene Ontology (GO) Biological Process enrichment analysis of differentially expressed genes.

**Supplementary Table S3.** WGCNA module membership (kME) values for all genes.

**Supplementary Table S4.** Nuclei counts and cell-type composition across samples, conditions, and iPSC lines.

**Supplementary Table S5.** Cluster-specific differential gene expression analysis following APAP treatment.

**Supplementary Table S6.** iPSC line information and sample metadata.

**Supplementary Table S7.** Experimental design summary for organoid lines, batches, and assays performed.

**Supplementary Table S8**. Source data and statistical analyses underlying the main figures.

